# Epistatic Transcription Factor Networks Differentially Modulate Arabidopsis Growth and Defense

**DOI:** 10.1101/583047

**Authors:** Baohua Li, Michelle Tang, Céline Caseys, Ayla Nelson, Marium Zhou, Xue Zhou, Siobhan M. Brady, Daniel J. Kliebenstein

## Abstract

Plants integrate internal and external signals to finely coordinate growth and defense allowing for maximal fitness within a complex environment. One common model for the relationship between growth and defense is a trade-off model in which there is a simple negative interaction between growth and defense theoretically driven by energy costs. However, there is a developing consensus that the coordination of growth and defense likely involves a more conditional and intricate connection. To explore how a transcription factor network may coordinate growth and defense, we used high-throughput phenotyping to measure growth and flowering in a set of single and pairwise mutants previously linked to the aliphatic glucosinolate defense pathway. Showing the link between growth and aliphatic glucosinolate defense, 17 of the 20 tested TFs significantly influence plant growth and/or flowering time. These effects were conditional upon the environment, age of the plant and more critically varied amongst the phenotypes when using the same genotype. The phenotypic effects of the TF mutants on SC GLS accumulation and on growth did not display a simple correlation, supporting the coordination model. We propose that large transcription factor networks create a system to integrate internal and external signals and separately modulate growth and the accumulation of the defensive aliphatic GLS.

## Introduction

Growth and defense are essential biological processes necessary for plant survival. Optimizing fitness requires plants to coordinate growth and defense phenotypes in response to specific environments. Efforts to understand the relationship between plant growth and defense is often modeled as a trade-off where resistance is a cost on growth (Karban and Baldwin 1997; Agrawal 2011a; Agrawal 2011b; Huot *et al.* 2014). This model assumes that the available resources for plants are limited, suggesting that any flux of resources, energy and elements into plant defense would be at the cost of lost plant growth. Support for this model comes from the observations that some constitutive defense mutants generally grow smaller and suffer from yield and/or fitness losses.

A developing model is emerging from research in ecology, evolutional biology and molecular genetics that suggests a dynamic relationship between plant defense and growth (Singh *et al.* 2002; de Lucas *et al.* 2016). Multiple reports showed that defense metabolism does not show a universal negative correlation with plant growth (Bergelson and Purrington 1996; Almeida-Cortez *et al.* 1999; Koricheva 2002). Mechanistic manipulations of defense metabolism have also shown little to no effect on plant growth. Further, diminutive constitutive defense mutants can have their growth rescued by second site mutations that maintain the constitutive defense (Hemm *et al.* 2003; Paul-Victor *et al.* 2010; Züst *et al.* 2011; Joseph *et al.* 2013; Campos *et al.* 2016; Kliebenstein 2016). Specific TFs are also being identified that can simultaneously increase growth and specific defenses (Campos *et al.* 2016; Wang *et al.* 2018). Together, these observations suggest that the relationship between plant defense and growth is a complex and active internal decision process involving regulatory and signaling pathways *in planta* (Zust and Agrawal 2017). Systemic and large scale studies are needed to fully explore the relationship between plant growth and defense, ideally in a highly characterized model system for both plant growth and defense.

Both plant growth and defense are regulated by complex signaling pathways, in which transcriptional factors (TFs) play key roles in integrating and transducing internal and external signals (Singh *et al.* 2002; Pajerowska-Mukhtar *et al.* 2012; Lozano-Duran *et al.* 2013; Fan *et al.* 2014; de Lucas *et al.* 2016). Several recent studies provide important insights about the roles TFs played in coordinating growth and defenses. Mutations of *JAZ*s, transcriptional repressors in jasmonic acid (JA) signaling pathway, in combination with altered photoreceptor *PhyB* results in fast-growing plants with enhanced plant defense responses (Campos *et al.* 2016). A rice TF, *Ideal Plant Architecture 1 (IPA)*, was previously cloned and characterized to activate yield-related genes and promote the high yield (Jiao *et al.* 2010), and recently further shown to promote both plant defense and rice yield simultaneously (Wang *et al.* 2018). These case studies demonstrated that growth and defense are under complex regulation and TFs are key integrators that coordinate these two important biological processes. However, it is not clear if these are isolated instances or a more generalizable view of how TFs modulate growth and defense. Systemic studies with well selected and designed experiments on TFs and their regulatory networks are needed to test how TFs may or may not coordinate between plant defense and growth in diverse environmental settings.

To explore the interplay between plant growth and defense, we used the classical and well-studied plant secondary metabolic pathway, the Arabidopsis methionine-derived aliphatic glucosinolates (GLS) (Sønderby *et al.* 2010b). The aliphatic GLS have been shown to provide defense against numerous herbivorous insects in both the lab and the field (Lambrix *et al.* 2001; Schlaeppi *et al.* 2008; Kos *et al.* 2012; Beran *et al.* 2014; Falk *et al.* 2014; Kerwin *et al.* 2015; Kerwin *et al.* 2017; Zalucki *et al.* 2017). Aliphatic GLS are critical for fitness in Brassicales and their specific composition and accumulation across developmental stages are intricately controlled by genetic variation and influence many important biological processes (Kliebenstein *et al.* 2002; Bidart-Bouzat and Kliebenstein 2008; Wentzell and Kliebenstein 2008; Burow *et al.* 2010; Kerwin *et al.* 2011; Züst *et al.* 2011). Thus, the aliphatic GLS are a key defense system within the Brassicales. Flux-based modeling suggests aliphatic GLS should be expensive to produce since they contain both sulfur and nitrogen and accumulate to high levels. However, mutants missing these compounds have at most a slight change in early growth, indicating that aliphatic GLS do not display a tight trade-off with growth (Paul-Victor *et al.* 2010; Züst *et al.* 2011; Bekaert *et al.* 2012). Instead, the regulatory complexity of the aliphatic GLS better fits with the coordination model. For example, a large scale yeast one-hybrid study identified a large collection of TFs that regulate aliphatic GLS accumulation (Li *et al.* 2014). These TFs bind the aliphatic GLS enzyme promoters, directly arguing against a single dominant TF model. Further, these TFs showed extensive epistatic interactions in influencing the accumulation of GLS (Li *et al.* 2018). This suggests that the regulation of this defense pathway is highly complex. However, previous studies did not inform on whether these TFs linked to aliphatic GLS have individual or epistatic effects on growth or flowering.

To test if growth and defense are linked by this epistatic network of TFs, we used a high throughput method to measure growth and flowering time of 20 single TF mutants and 48 double mutants previously shown to influence the Arabidopsis aliphatic GLS pathway in two environments (Sønderby *et al.* 2007; Li *et al.* 2018). We show that 17 of the 20 TFs significantly influence plant growth and/or plant flowering time. While most TFs influence growth and defense, there was no clear correlation between the aliphatic short chain (SC) GLS accumulation and plant growth and flowering time in our study. This indicates that each TF has specific and independent influences on both growth and aliphatic GLS. Our findings support the coordination model for the relationship between plant growth and aliphatic GLS, and provide novel insights on how these critical and complex biological processes are integrated to optimize fitness in different environments.

## Results

### Conditional Growth Effects of the 20 TF Mutations

The 20 selected TFs for this study were originally identified as binding aliphatic GLS related promoters and their knockout mutants influence aliphatic GLS accumulation (Li *et al.* 2014; Li *et al.* 2018). In this study, we focused on the aliphatic GLS as they are ∼90% of the total GLS content within Arabidopsis leaves representing the majority of the metabolic flux. To measure growth variation in this mutant collection which have altered defense levels that are within the range of variation occurring within natural accessions (Kliebenstein *et al.* 2001), we utilized digital image analysis to measure plant size in two chambers, CEF clean chamber and LSA herbivore chamber (Li *et al.* 2018). Growth was measured every other day from 9 days post-germination to 27 days at which time the leaves were overlapping. Plants were organized in a randomized complete block design with 8 measurements per genotype per chamber (Li *et al.* 2018). The two growth chambers differed in the presence or absence of mites, flea beetles and fungus gnat larvae, allowing us to test how the underlying networks respond to complex environmental perturbations. To maintain this difference, CEF was cleaned monthly while LSA had long term tomato and Brassica plants that sustained the endogenous pest population. The plant growth measurement spans the majority of vegetative growth providing a dynamic analysis of growth. Combining the data with linear models, we tested for significant effects of all single gene TF mutants on growth across the conditions (Figure 1, Supplemental Figure 1 and Supplemental Data Set 1, 2). To reduce false positives, we required a TF to have at least 2 significant hits across the days in the TF and/or treatment conditioned TF effect to be validated as TF influencing plant growth.

**Figure 1.**
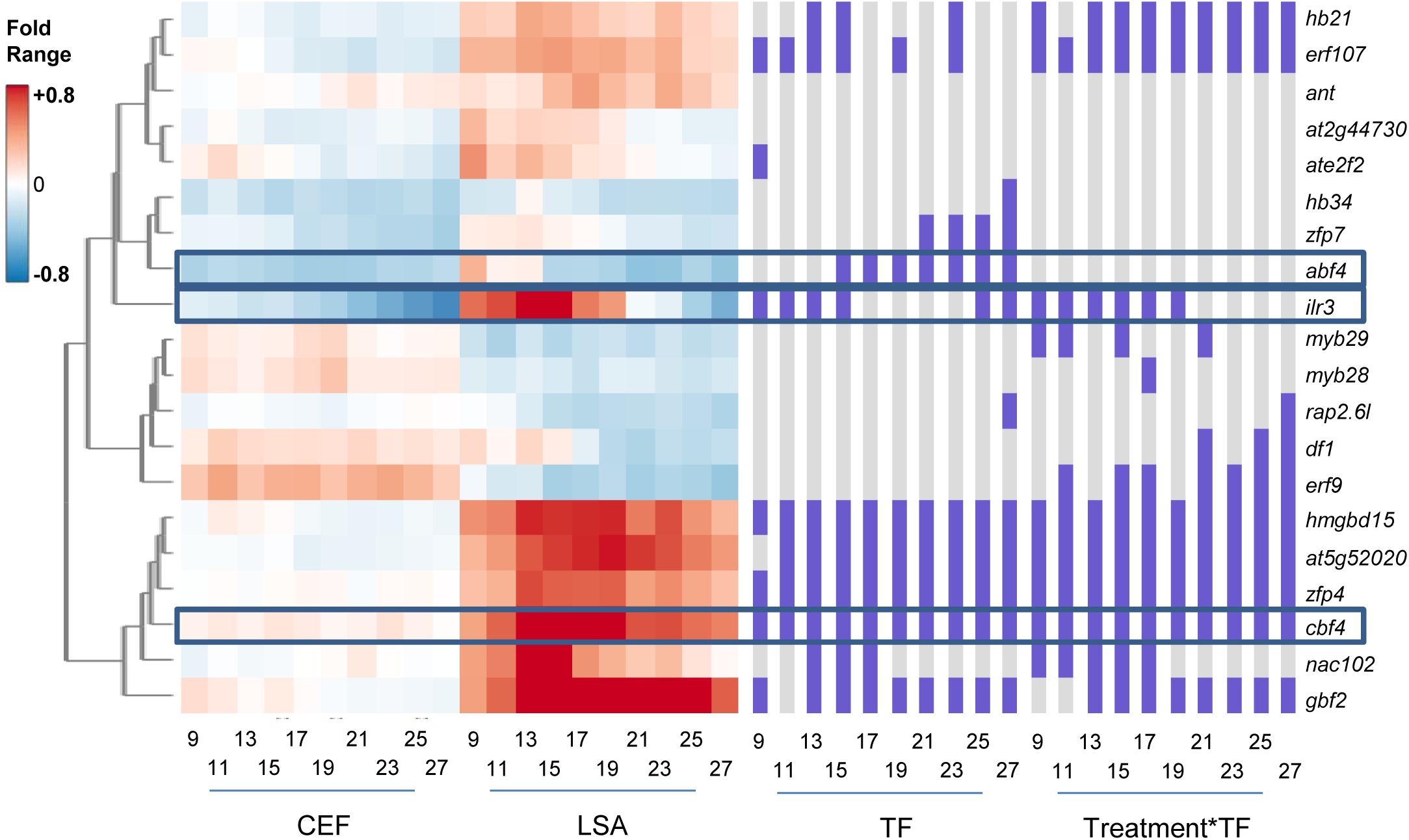
The Effects of the 20 TFs on Plant Growt The heatmap displays the fold change of plant size in single mutants, from day 9 to day 27, in comparison to Col-0 in both the clean CEF chamber and herbivore LSA chamber. Red shows increased plant size and blue decreased plant size. The columns on the right display the statistical significance (purple, significant p< 0.05; gray, not significant) for each term in the statistical model as listed at the bottom. Specific framed genotypes have mean phenotypes shown for reference in Supplemental Figure 1.

The influence of these TFs on growth is highly conditional on both plant age and growth chamber (Figure 1 and Supplemental Figure 2). 15 of the 20 aliphatic GLS TF mutants had a significant influence on growth. Stronger positive growth effects were observed more often in the herbivore LSA chamber than in the clean CEF chamber for a number of TFs like *HMGBD15, AT5G52020, ZFP4, CBF4, NAC102*, and *GBF2* (Figure 1). In contrast, *ABF4* had strong negative growth effects in both chambers. In addition to differences between the growth chambers, there were TFs that had differential influence across plant age. For example, *GBF2* has growth effects across most plant stages while *nac102* only shows a significant phenotype during early vegetative growth (Figure 1). Some mutants combined these conditionality’s as illustrated by *ilr3*, which had a transition at 21 days post germination from positive to negative growth effects. This conditionality illustrates the importance of large-scale phenotyping across different conditions to generate a broad view of how mutational effects may change dynamically (Supplemental Figure 1 and 2). In an energetic tradeoff, we would expect that most of these TF mutations should lead to smaller plants as these mutations increase aliphatic GLS content (Li *et al.* 2014). In contrast, the mutants had a mix of positive and negative growth effects. In addition, three mutants with strong aliphatic GLS phenotypes, *ant, myb28* and *myb29*, have little to no detectable main effect on growth. Although an alternative explanation for the *myb28/29* mutant is that this mutation has a ∼2x increase in indolic GLS accumulation and this may balance the loss in aliphatic GLS. However, because aliphatic GLS are ∼90% of the total GLS, this slight increase in indolic GLS means that the *myb28/29* double mutant has a loss of 80% of total GLS flux. The links between plant growth and defense via the aliphatic GLS in this TF collection supports the coordination model of growth and defense.

### Dynamic Epistatic Networks Underlying Plant Growth

We previously generated a set of 48 pairwise mutant combinations from these 20 TFs that identified an extensive antagonistic epistatic network of the 20 TFs controlling aliphatic GLS (Sønderby *et al.* 2007; Li *et al.* 2018). As these TFs had affected growth individually, we used linear models to assess if the epistatic effects were as prevalent on growth as on aliphatic GLS accumulation. We plotted the significant interactions on growth for five different days on the aliphatic GLS epistatic network (Figure 2A-E, and Supplemental Data Set 3, 4). This analysis identified extensive epistatic interactions influencing growth. As observed in the single mutants, the epistatic interactions were equally conditional across ontogeny and growth condition (Figures 2 and 3). Interestingly, there were more epistatic interactions identified at Day 17 than at earlier or later growth stages. This is possibly because some interactions were present during earlier developmental stages while others were present during later developmental stages. Day 17 was the age at which early and late patterns appeared to overlap. This suggests a transition in the epistatic network and underlying mechanisms around this time. Interestingly, the strong aliphatic GLS TFs, *MYB28, MYB29, ANT*, that had minimal single mutant influence on growth, had significant epistatic interactions on plant growth (Figure 2 and 3). Thus, this set of TFs identifies and illustrated an epistatic system influences plant growth depending on age and growth condition. The effects of these TFs on growth are more conditional than the aliphatic GLS effects suggesting that while the traits of growth and defense are both controlled by the epistatic network, they are independent regulatory outputs.

**Figure 2.**
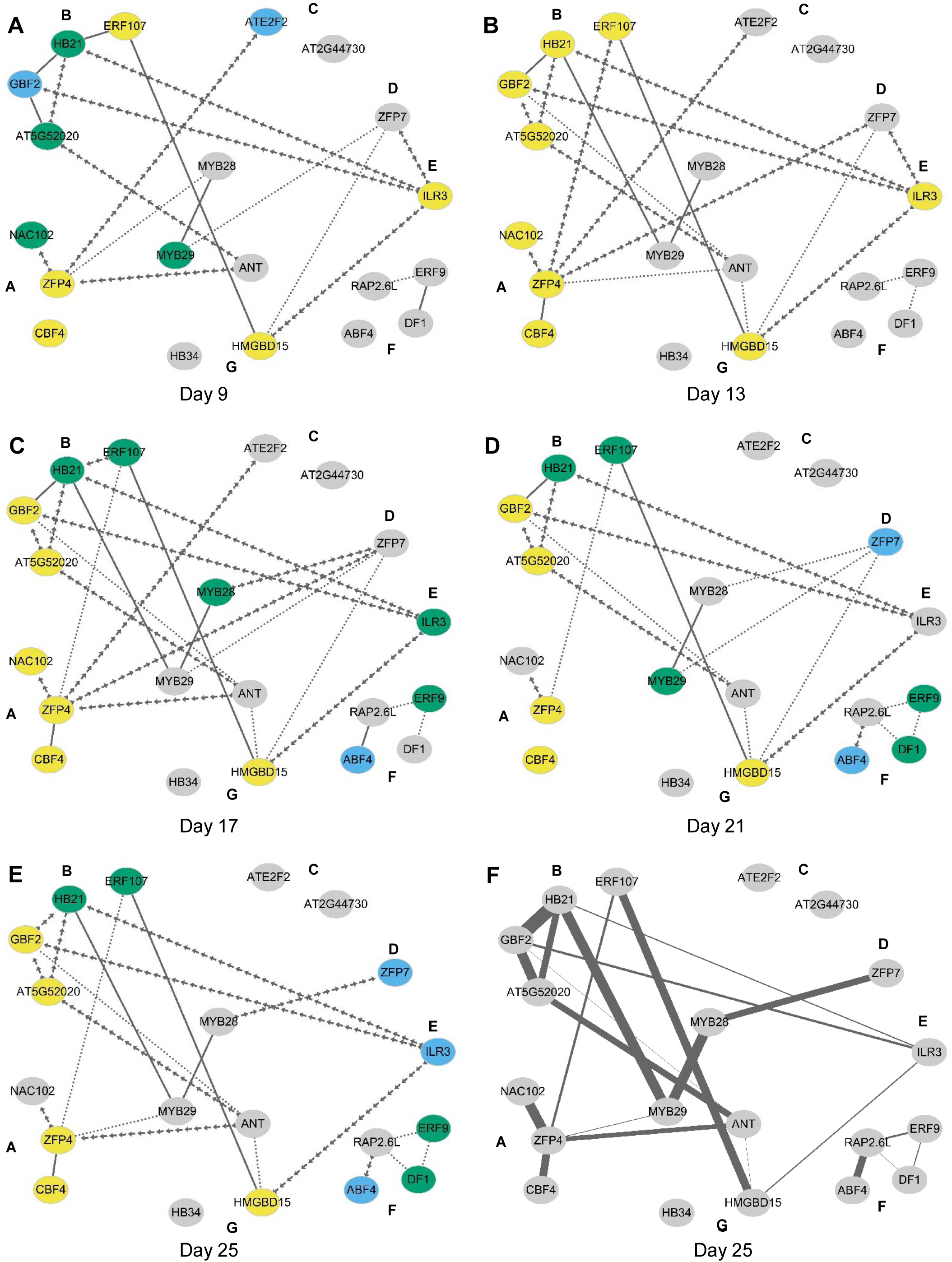
Epistatic networks for representative growth days Connectivity plots of epistatic interactions between TF genes are shown. The TFs are laid out in the network based on their effect on aliphatic GLS accumulation. Solid lines show that the TF x TF interaction term was significant while dotted shows significant Treatment x TF x TF interaction in the statistical model. A line of arrows shows that the interaction was conditional on both treatment and tissue. The color of the node indicates which main effect terms are significant for the individual TFs; sky blue indicates only a TF main effect, green indicates only a treatment x TF interaction, yellow indicates both TF and treatment x TF. A-E: Shows the network significances on Day 9, 13, 17, 21, and 25. F: Visualization of individual epistatic variance components within the genetic network for day25. The width of the line connecting 2 TFs is proportional to the variance linked with the TF x TF term for that specific interaction.

### Conditional Epistasis Networks Underlie Plant Growth

To quantify the epistatic effects on plant growth, we used a previously established epistasis value to measure the direction and magnitude of each epistatic interaction (Li *et al.* 2018). Briefly, we subtracted the measured double mutant phenotype from the predicted double mutant phenotype under an additive model. This value was then normalized to the wild type phenotype. This epistasis value was measured for each pair of mutants for all the growth data (Supplemental Data Set 5). The epistasis value will be positive when there is synergistic epistasis and negative for antagonistic epistasis (Figure 3). Previous work showed that epistasis for the short chain GLS (SC GLS), the dominant form of aliphatic GLS, was almost entirely negative/antagonistic (Li *et al.* 2018). Unlike SC GLS, epistasis for growth was a mix of antagonistic and synergistic values that could shift depending upon the conditions and developmental stages. For example, *rap2.6l* was involved in several epistatic interactions that were positive in the LSA growth condition but negative in the CEF growth condition (i.e. *rap2.6/erf9*, etc.); in contrast, epistatic interactions involving *hmgbd15* had negative interactions in the LSA and positive in the CEF growth conditions (i.e. *ilr3/hmgbd15*); other TF combinations like *myb28/myb29* had similar epistasis across both growth conditions (Figure 4). This argues that this epistatic network of TFs influences growth in both conditions but that the environmental signals in the two conditions permeate differently through the TF network to generate variable growth outputs.

**Figure 3.**
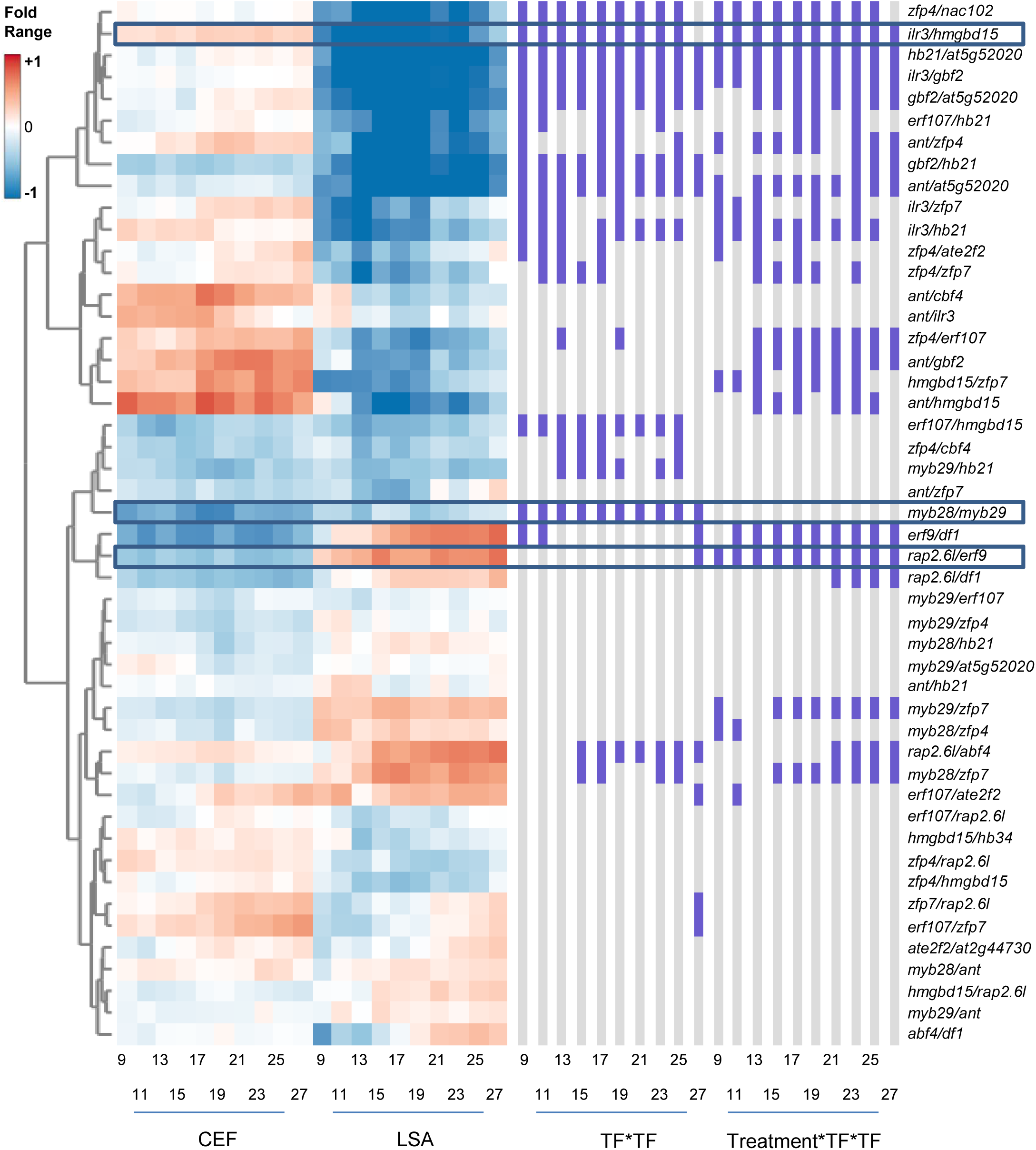
Epistatic effects on plant growth Epistasis values were calculated for all pairwise TF combinations from day 9 to day 27 in each treatment condition. These values are plotted in a heatmap for all pairwise mutant combinations. The genotypes are clustered using hierarchical clustering and labeled to the right of the diagram. The columns to the right of the heat map show which epistatic interaction term is significant (purple) or not significant (gray) (P < 0.05). Specific framed genotypes have mean phenotypes shown for reference in Figure 4.

**Figure 4.**
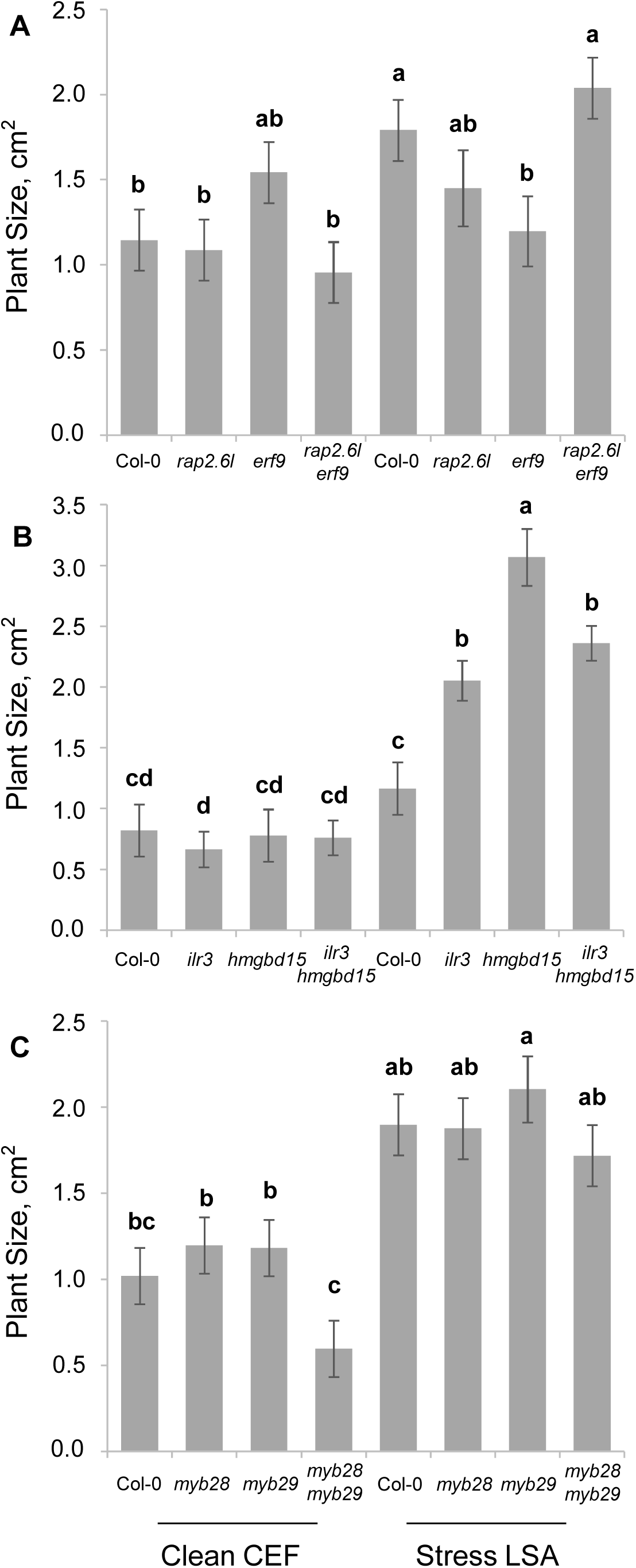
Representative epistatic growth phenotypes The average rosette size on day 15 for the corresponding genotypes from the clean CEF and herbivore LSA chambers are plotted. Different letters indicate genotypes with significantly different plant sizes. (P < 0.05 using post-hoc Tukey’s test). SE is shown with 8 samples across two experiments for each genotype. Day 15 was chosen to provide a common date across which to illustrate key differences. (A) Rosette size of single and double mutants of *rap2.6l* and *erf9* (B) Rosette size single and double mutants of *ilr3* and *hmgbd15* (C) Rosette size single and double mutants of *myb28* and *myb29*

### Epistatic Networks Influence Flowering Independent of Tested Environments

To test if these TFs and their network influence reproduction, we measured flowering time in all of the plants from all of the genotypes in the two contrasting chambers, and could show that 12 of the 20 tested TFs significantly influenced flowering (Figure 5). However, unlike growth, the effects on flowering time were almost entirely towards early flowering in both chambers, with only *ANT, ILR3* and *ERF9* having an environmental conditionality (Figure 5, and supplemental figure 3). Plotting the epistatic effects for flowering time showed predominantly synergistic epistasis with 18 statistically significant interactions and only 2 interactions having environmental conditionality (Figure 6, A and B). This indicates that the genetic control of flowering time is less influenced by the environmental conditions than are either growth or aliphatic GLS. To further visualize how the epistatic variance was influenced by the network topology, we mapped the epistatic variance for the plant flowering (Figure 6C). Using previously ascribed groupings of the TFs based on their GLS phenotypes, TFs in Group A and B have more significant epistatic interactions partly overlapping with the epistatic interactions of growth phenotype, and TFs with strong main effects on flowering time also have higher genetic variance in their epistatic interactions.

**Figure 5.**
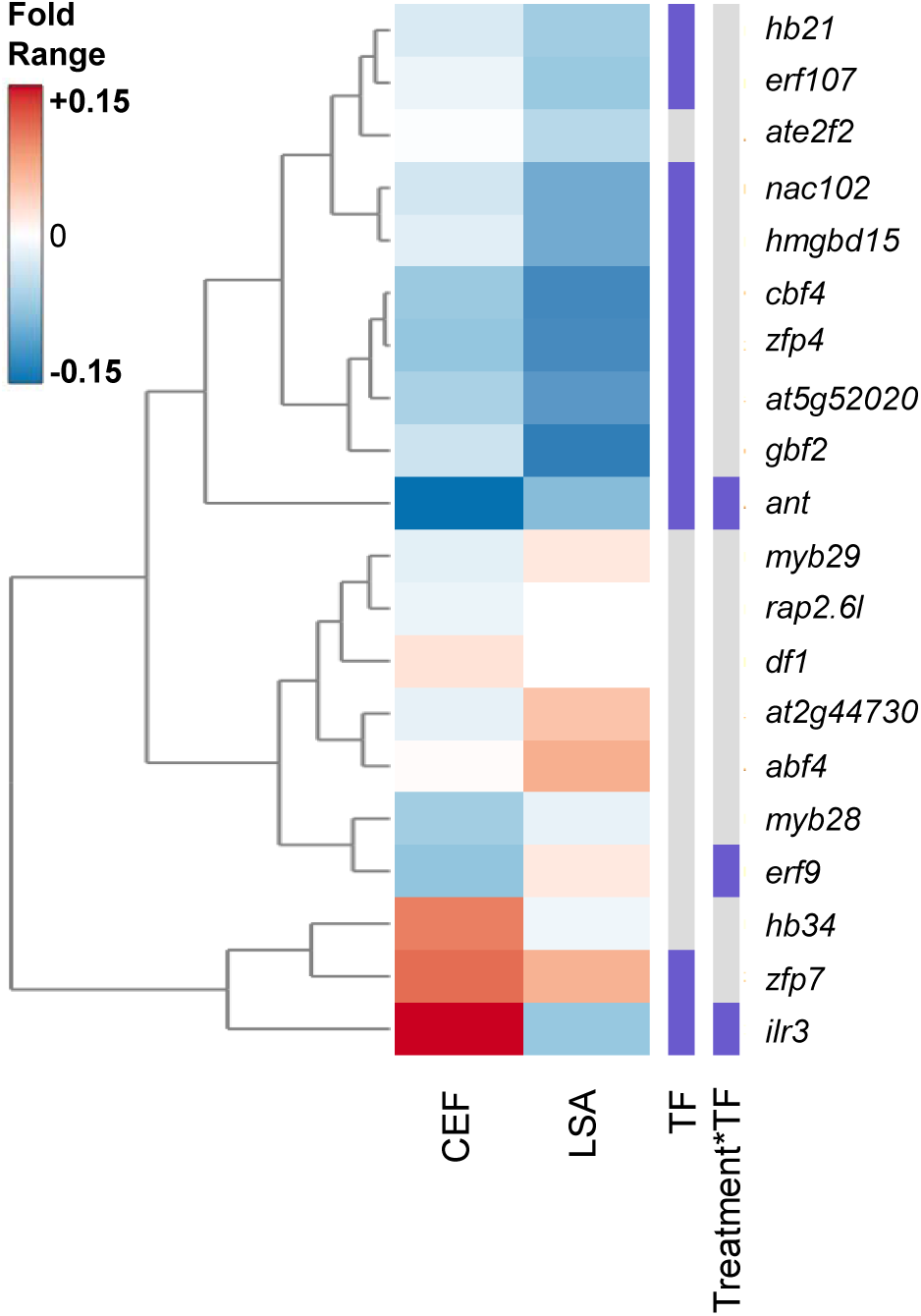
TF Effects on Flowering Time The heatmap displays the fold change of flowering time in the single mutants compared with Col-0 in both clean CEF chamber and herbivore LSA chamber. Red shows increased flowering time and blue decreased flowering time. The columns on the right display the statistical significance (purple, significant p< 0.05; gray, not significant) for each term in the statistical model as listed at the bottom.

**Figure 6.**
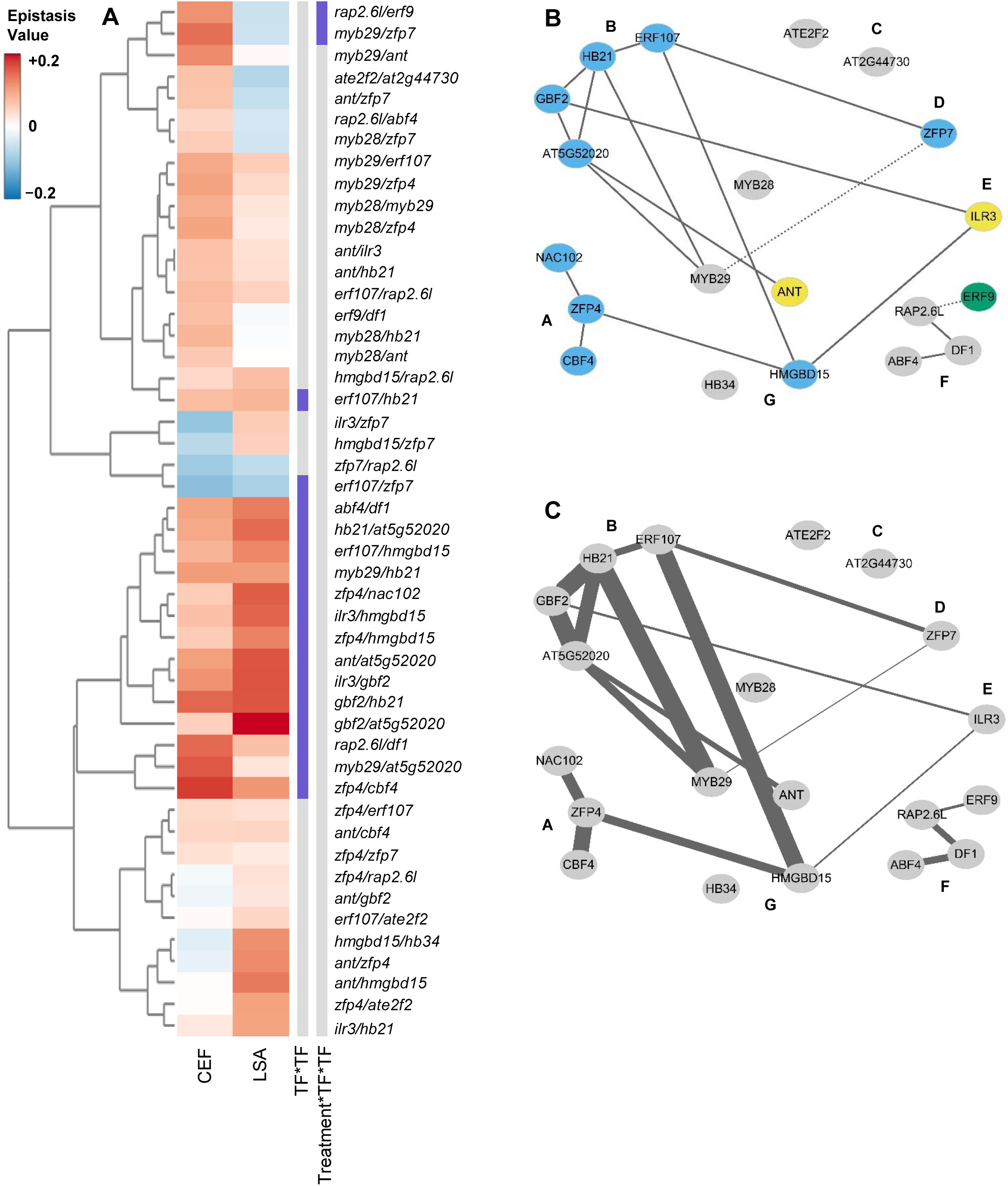
Epistatic network and flowering time effects A: Epistasis values were calculated for all pairwise combinations individually in both treatment conditions and plotted in the heatmap. The genotypes are clustered using hierarchical clustering and labeled to the right of the diagram. The columns to the right of the heatmap show which epistatic interaction term is significant (purple) or not significant (grey) (P < 0.05). B: A representation of the significant epistatic networks is shown for flowering time. Solid lines show that only the TF x TF interaction term was significant and a dotted line shows where there was a significant chamber x TF x TF interaction. The node color indicates which main effect terms are significant for the individual TFs; sky blue indicates only a TF main effect, green indicates only a chamber x TF interaction, yellow indicates both TF and chamber x TF. C: Visualization of individual epistatic variance components within the genetic network for flowering time. The width of the line connecting 2 TFs is proportional to the variance linked with the TF x TF term for that specific interaction.

### Connections between Defense and Growth

To further explore the connection between plant defense and development, we systemically tested for associations between growth and defense in this data. These TFs predominantly influence the accumulation of SC GLS, and we used this as our quantification for defense (supplemental data set 1) (Li *et al.* 2018). The mutant effects of the 20 TFs on SC GLS, plant growth and flowering time in both chambers were calculated by taking the phenotypic difference between mutant and wild-type and normalized by the wild-type value. We then tested for a relationship between growth and defense by testing for correlations between the mutant effects using these traits (Figure 7, Supplemental data set 6). In contrast to the expectation if this system was being solely driven by an energetic trade-off model, the TFs’ effects on the accumulation of SC GLS and growth/flowering were largely uncorrelated (Figure 7A and B). This was equally true in both the stress and the clean chamber, indicating that the absence of biotic attackers did not illuminate a hidden relationship. As expected, there was a correlation between growth and flowering. Our findings show a highly complex relationship between plant defense and plant growth.

**Figure 7.**
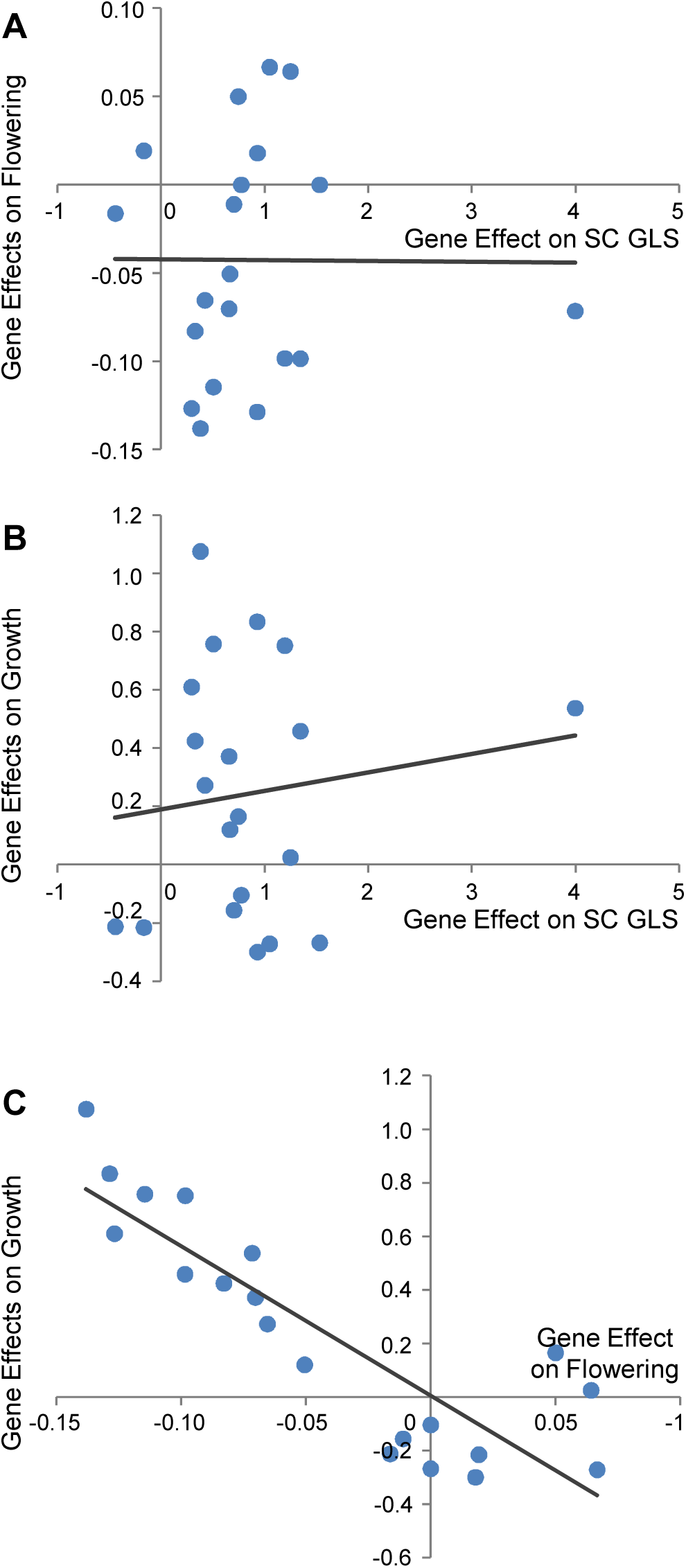
Lack of relationship between mutant effects on defense and growth. The main additive effect of the TF mutants in herbivore LSA chamber were selected to illustrate the absence of a relationship between growth, flowering time and SC GLS. The predicted linear tread line and the statistical test results are shown. A: Comparison of gene effects on SC GLS and flowering time (r = −0.006, p = 0.980). This result was unaffected by the presence or absence of the outlier point for SC GLS. B: Comparison of gene effects on SC GLS and growth (r = 0.131, p = 0.583). This result was unaffected by the presence or absence of the outlier point for SC GLS. C: Comparison of gene effects on flowering time and growth (r = −0.867, p < 0.001).

### The Connection of Single Mutant and Double Mutant Effects

Given the low correlation between the SC GLS accumulation and plant growth, we proceeded to investigate if there may be any connection between a TF’s main effect on a trait and its average epistatic effect on the trait. This allows us to investigate if there is any internal influence of single gene effects on the direction and value of the epistatic value from their double mutants. If there is significant correlation, it would further advance our understanding of internal properties of the epistatic network and revealed underlying quantitative genetic principle governing all the phenotypic changes. To do this, we calculated the average gene epistasis value for each trait for each single TF across all the pairwise combinations. Next, we correlated the TF’s estimated single mutant effects to their average gene epistasis value for each phenotype and chamber combinations (Figure 8, Supplemental data set 6). The analysis showed that within all 3 tested traits, there are strong, consistent and negative correlations between a mutation’s main and epistatic effects. One possible interpretation of this result is that the genetic background is constraining the results that we have obtained. The prevalence of negative epistasis in the SC GLS phenotype agrees with this possibility. Future work is required to understand if this observation is a general property of this network or is a function of the specific Col-0 accession in which it was conducted.

**Figure 8.**
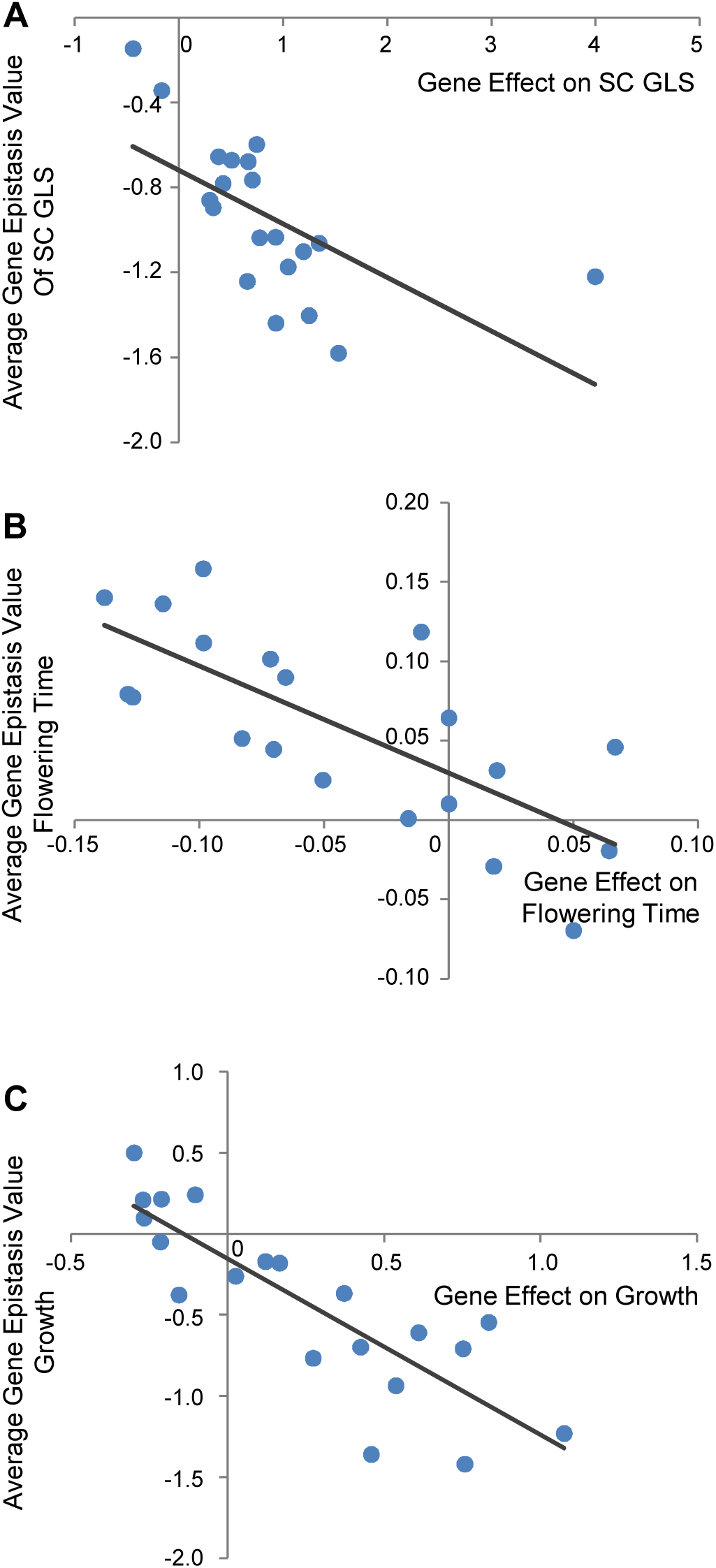
Relationship of the main and epistatic effects for TF genes The average epistatic effect of a TF across all its pairs in the herbivore LSA chamber was calculated and compared to the TFs main effects for SC GLS, flowering time and plant size on Day 17. The predicted linear tread line and the statistical test results are shown. A: Comparison of main gene and average epistatic gene effects on SC GLS (r = −0.610, p = 0.004). B: Comparison of main gene and average epistatic gene effects on flowering time (r = −0.732, p < 0.001). C: Comparison of main gene and average epistatic gene effects on flowering time on growth (r = −0.841, p < 0.001).

## Discussion

In this study, we tested 20 TF mutants and their 48 double mutants that influence aliphatic GLS accumulation to systemically explore if and how these TFs could significantly influence plant growth and flowering time. 17 of these 20 TFs significantly influence both plant growth and flowering time. Interestingly, the key aliphatic GLS regulators *MYB28* and *MYB29* have little influence on plant growth and flowering, and what is more, there is no simple mechanistic connection between the TFs’ effects on the accumulation of aliphatic SC GLS and growth or flowering. These findings fit with the emerging coordination model whereby the network is dynamic and highly responsive to the specific requirements of a specific environment. We propose that the internal and external signals are coordinated and perceived via connections between these diverse TFs. These connections create a decision matrix whereby growth and aliphatic GLS are interpreted and coordinated. Independent connections to growth and aliphatic GLS then proceed from this matrix to maximize plant fitness in a given environmental setting. (Figure 9). Further, this suggests that TF networks provide an unappreciated potential to fine-tune both growth and defense to optimize modern agriculture.

**Figure 9.**
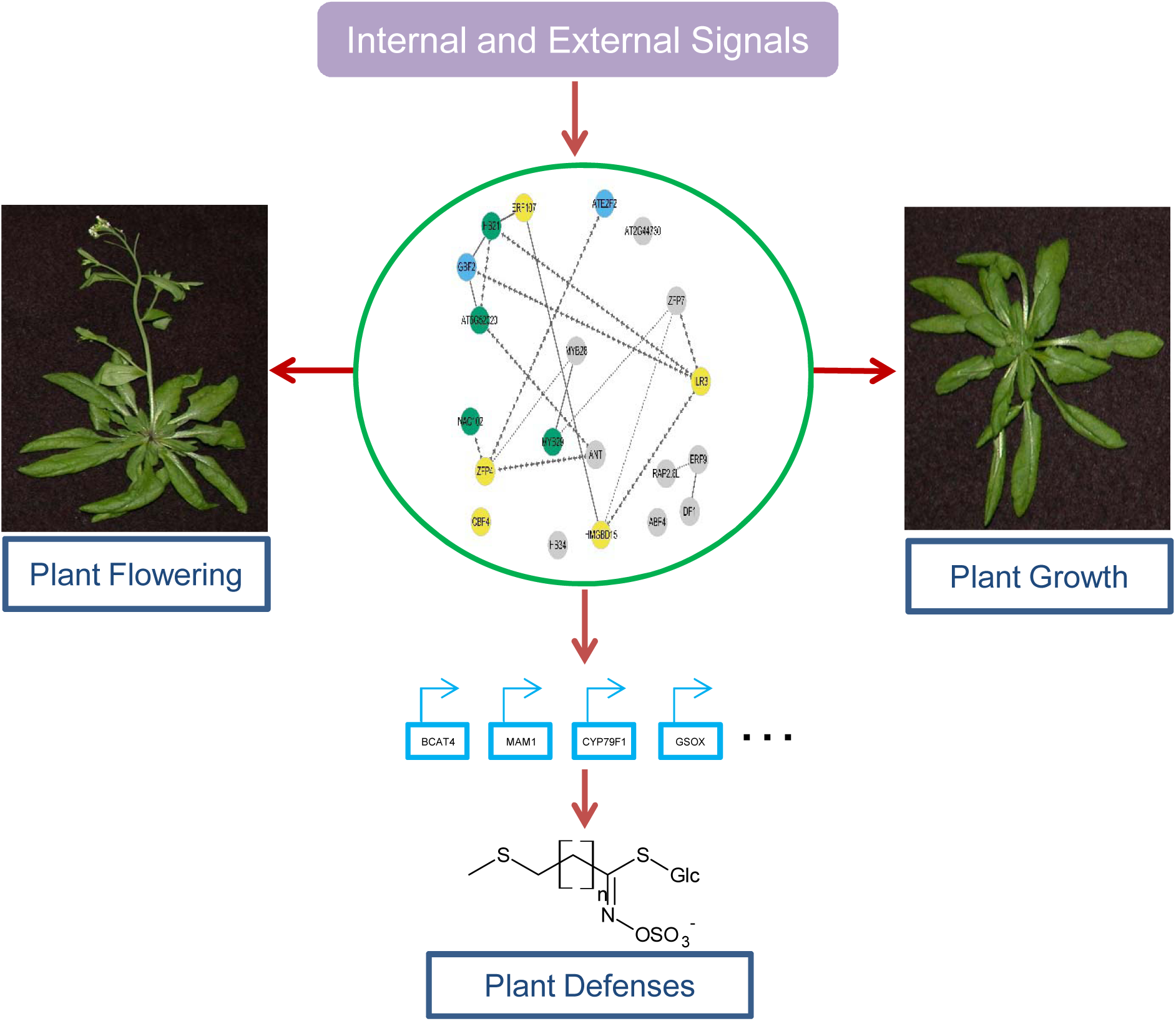
Proposed model for TF coordination of plant development and aliphatic GLS based plant defense External environmental and internal developmental signals are coordinately perceived via a group of TFs. Direct and indirect connections between these TFs allow for a coordinated response to these complex signals. A theoretically cohesive response is then transmitted to growth and defense via separate outputs from this network.

### Growth and Defense vs Coordination

The canonical model for plant growth and plant defense is a trade-off model, which treats plant growth and defense as two competing biological processes under the assumption that the acquisition of elements and energy are limiting, a zero sum game. A complementary hypothesis is that growth and defense are two separable outputs of the plant’s regulatory system that must be coordinated depending on the specific environment. The analysis of the phenotypes in this TF collection and epistatic network in aliphatic GLS pathway supported the existence of the coordination model. The complex interactions between growth and defense depended on the specific perturbed TF, the environment and their epistatic effects. There was no consistent evidence of any negative relationship between the accumulation of the SC GLS defense metabolites and any measurement of growth. This suggests that the TFs collectively make growth and defense decisions in a coordinated fashion in our case study, and the final growth and defense phenotypes are the outcomes from this decision process. Future work is required to test if this decision process is truly structured to shape both growth and defense in a way that always maximizes the individual’s fitness potential in their ecological niche. This suggests that future experiments need to incorporate more complex regulatory relationships to truly understand how growth and defense are related in the field.

### Using Transcription Factors to Tune and Optimize Growth and Defense

Putative costs of plant defenses on growth have been intensively researched, and recent findings are suggesting that it is possible to promote one without sacrificing the other (Campos *et al.* 2016; Wang *et al.* 2018). Our current study further brightens the future potential to optimize plant growth and defense through TF manipulation. Specifically, we found that there are a vast array of potential TFs that can be identified and potentially used to manipulate plant growth and defense. In our system, mutations in more than half of the tested TFs could significantly promote plant growth and defense together across diverse environmental conditions. Critically, this ability to promote both defense and growth is highly conditional on the specific environment in which the phenotypes are being measured. If this is true in other genes and TFs involving plant growth and defense, it raises the importance of studying plants’ response to genetic manipulation under different growth conditions, which is especially relevant and pertinent to crop breeding efforts in changing climate and environmental stresses. To fully interrogate this potential, future studies are needed that involve larger collections of TFs and epistatic interactions across even more diverse environments to further understand the coordination between growth and defense across fluctuating environments.

## MATERIALS AND METHODS

### Plant materials and growth conditions

The *Arabidopsis thaliana* T-DNA insertion lines of the 20 TFs were ordered from Arabidopsis Biological Resource Center (Sussman *et al.* 2000; Alonso *et al.* 2003) and homozygous lines were validated in previous studies (Sønderby *et al.* 2010a; Li *et al.* 2014). The 47 double mutants were generated and validated and planted in previous study (Li *et al.* 2018). Briefly, the Arabidopsis plants were grown in two independent chambers with 16-h light at 100- to 120-µEi light intensity with temperature set at a continuous 22°C. The two growth chambers were set to similar abiotic environments but contain dramatically different biotic environments, one pest free, clean CEF as a typical lab growth condition, and one with an endogenous pest population, herbivore LSA mimicking the nature growth condition. Seeds were imbibed in water at 4°C for 3 days and sown into Sunshine Mix (Sun Gro Horticulture). Seedlings were thinned to one plant per pot (6cm × 5cm) at 7 d after planting. For each experiment, at least 8 replicates of Col-0, single and double were planted using a randomized complete block design. Each flat had one plant per genotype leading to eight flats per replication. This experiment was conducted independently in the clean CEF and herbivore LSA chamber to generate a minimum of 16 biological repeats in total for most of the genotypes.

### Flowering time and plant growth measurement

All the plants were checked daily, and the day was recorded for each of the plants when the first flower opened. The flowering time was measured as how many days it takes for the plant to have the first flower opened. All the plants were taken photos from day 9 to day 27 when some of the early flowering lines in stress chamber started to flower. The circling area of the rosette of each of the plant in each growth condition was manually measured by Image J across the days. The area was used as an indicator of the plant size. The plant flowering time and plant sizes were normalized with the number of Col-0, and further visualized using the iheatmapr package in R software (R Development Core Team 2014).

### Statistics

To test for epistasis of the TFs in controlling plant development, the flowering time and plant growth for each epistatic combination were separately analyzed by ANOVA using a general linear model with lmerTest in R (Kuznetsova *et al.* 2017). The following model was used to test for the epistasis for the flowering time and growth phenotypes in the double mutants, with each double mutant having both single mutants and wild type grown concurrently: *y*_*abc*_ *= µ* + *A*_*a*_ +*B*_*b*_ + *Ch*_*c*_ + *A*_*a*_*xB*_*b*_ + *A*_*a*_*xCh*_*c*_+ *B*_*b*_*xCh*_*c*_ + *A*_*a*_*xB*_*b*_*xCh*_*c*_+ *ε*_*abc*_, where *ε*_*abc*_ is the error term and is assumed to be normally distributed with mean 0 and variance σ_ε_^2^. In this model, *y*_*abc*_ denotes the plant flowering and growth data in each plant, Genotype A represents the presence or absence of a T-DNA insert in one TF gene (WT versus mutant of locus A), and Genotype *B* represents the presence or absence of a T-DNA insert in another TF gene (WT versus mutant of locus B) in the double mutant from Chamber *Ch*_*c*_ (Clean CEF chamber or herbivore LSA Chamber). The ANOVA table, least-square (LS) means and standard error for each genotype x treatment combinations were obtained with emmeans package in R (Searle *et al.* 1980). The type III sums-of-squares from this model were used to calculate the variance and percent variance attributable to each term in the model. For the percent variance, this was calculated by comparing to the total variance in the model as the denominator. All network representations were generated using Cytoscape.v2.8.3 (Shannon *et al.* 2003).

### Calculation of epistasis value

To study the effect of epistasis, we use epistasis value to describe the direction and strength of the epistasis by normalizing the difference of observed double mutant phenotype versus the predicted double mutant phenotype, assuming additivity of the single mutants, then normalized to the wild type as reported before (Segre *et al.* 2005; Li *et al.* 2018).The phenotype for WT was set as w, mutant TFa as a, mutant TFb as b, and double mutant TFa/TFb as ab. The Epistasis Value is calculated as (ab - (w + (a-w) + (b-w))/w). If the epistasis value is positive, this shows evidence for synergistic epistasis, while antagonistic epistasis is reflected in negative values. The larger the epistasis value, the stronger the epistasis effects. The Epistasis Value were further visualized using the iheatmapr package in R software (R Development Core Team 2014).

## Supporting information

Supplemental Data Set 5

Supplemental Data Set 3

Supplemental Data Set 4

Supplemental Data Set 2

Supplemental Data Set 1

Supplemental Data Set 6

## Acknowledgements

Funding for this work was provided by the NSF awards IOS 1655810 and 1547796 to DJK, the USDA National Institute of Food and Agriculture, Hatch project number CA-D-PLS-7033-H to DJK and by the Danish National Research Foundation (DNRF99) grant to DJK. SMB was partially funded by an HHMI Faculty Scholar Fellowship.

---

Supplemental Figure 1. Representative plant growth phenotypes in single TF mutants Rosette sizes on day 15 for the corresponding genotypes are shown in the clean CEF and herbivore LSA chambers. Different letters indicate genotypes with significantly different plant sizes. (P < 0.05 using post-hoc Tukey’s test). SE is shown with 8 samples across two experiments for each genotype.

A. *cbf4*
B. *abf4*
C. *ilr3*

Supplemental Figure 2. Dynamic Mutant Effects on Plant Growth

Rosette growth of specific mutations in comparison to Col-0 is shown across time. Col-0 is shown as a solid line while mutants are the dashed line. Data from the CEF clean chamber are shown as black lines data from the herbivore LSA chamber are red lines.

A. *cbf4*
B. *abf4*
C. *ilr3*

Supplemental Figure 3. Representative flowering time phenotypes in single TF mutants Flowering time in specific genotypes is shown for both the clean CEF and herbivore LSA chamber. Different letters indicate genotypes with significantly different flowering time. (P < 0.05 using post-hoc Tukey’s test). SE is shown with 16 samples across two experiments for each genotype.

A. *zfp7*
B. *ant*
C. *ilr3*

