## Supplemental Figures for "Epistatic Transcription Factor Networks Differentially Modulate Arabidopsis Growth and Defense"

### Slide 1
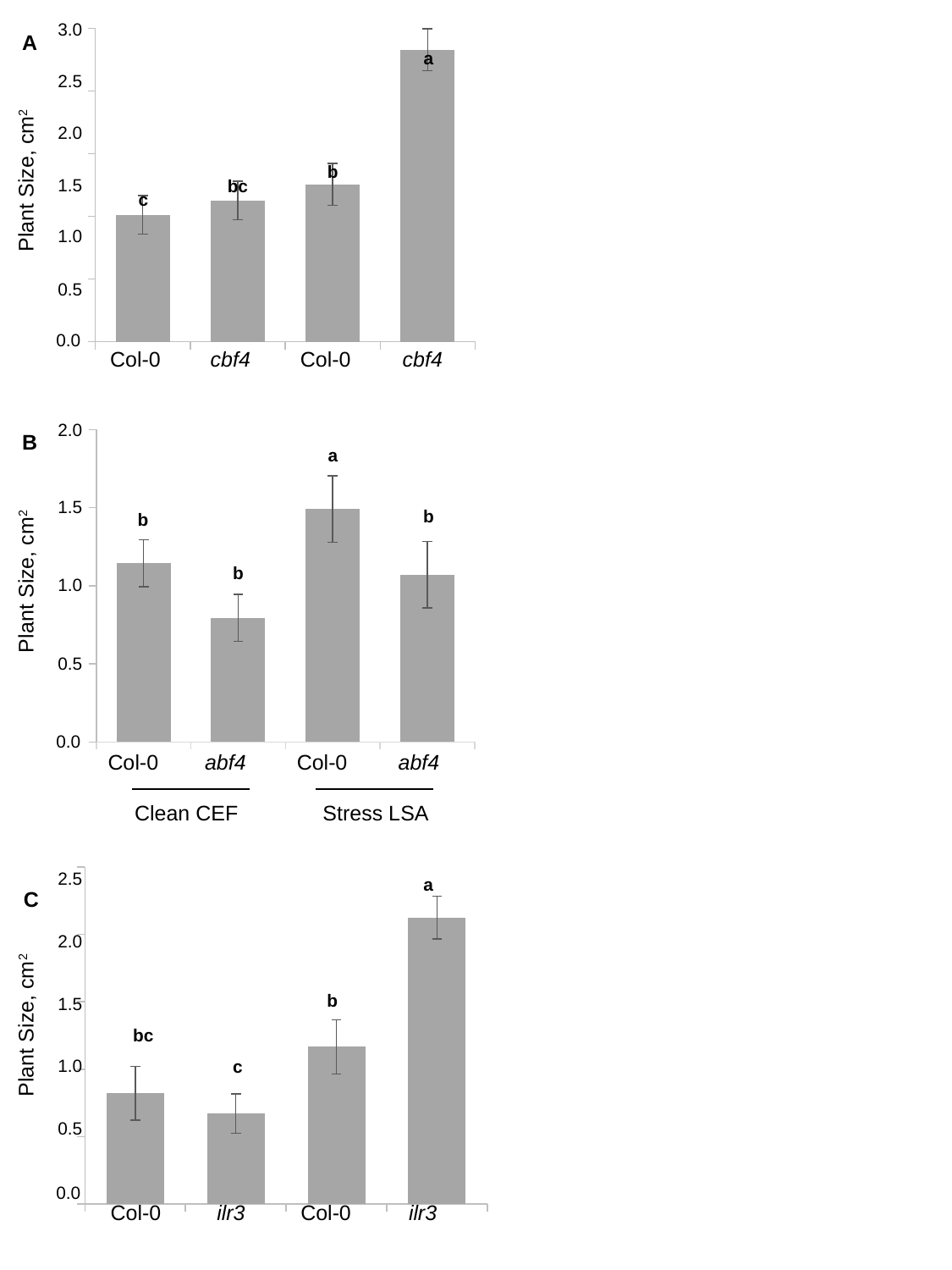

#### Chart
| Category | |
|---|---|3.0
A
a
2.5
Plant Size, cm^2^
2.0
b
1.5
bc
c
1.0
0.5
0.0
Col-0
Col-0
cbf4
cbf4
#### Chart
| Category | |
|---|---|2.0
B
a
1.5
b
b
Plant Size, cm^2^
b
1.0
0.5
0.0
Col-0
Col-0
abf4
abf4
Clean CEF
Stress LSA
#### Chart
| Category | |
|---|---|2.5
a
C
2.0
Plant Size, cm^2^
b
1.5
bc
1.0
c
0.5
0.0
Col-0
Col-0
ilr3
ilr3

### Slide 2
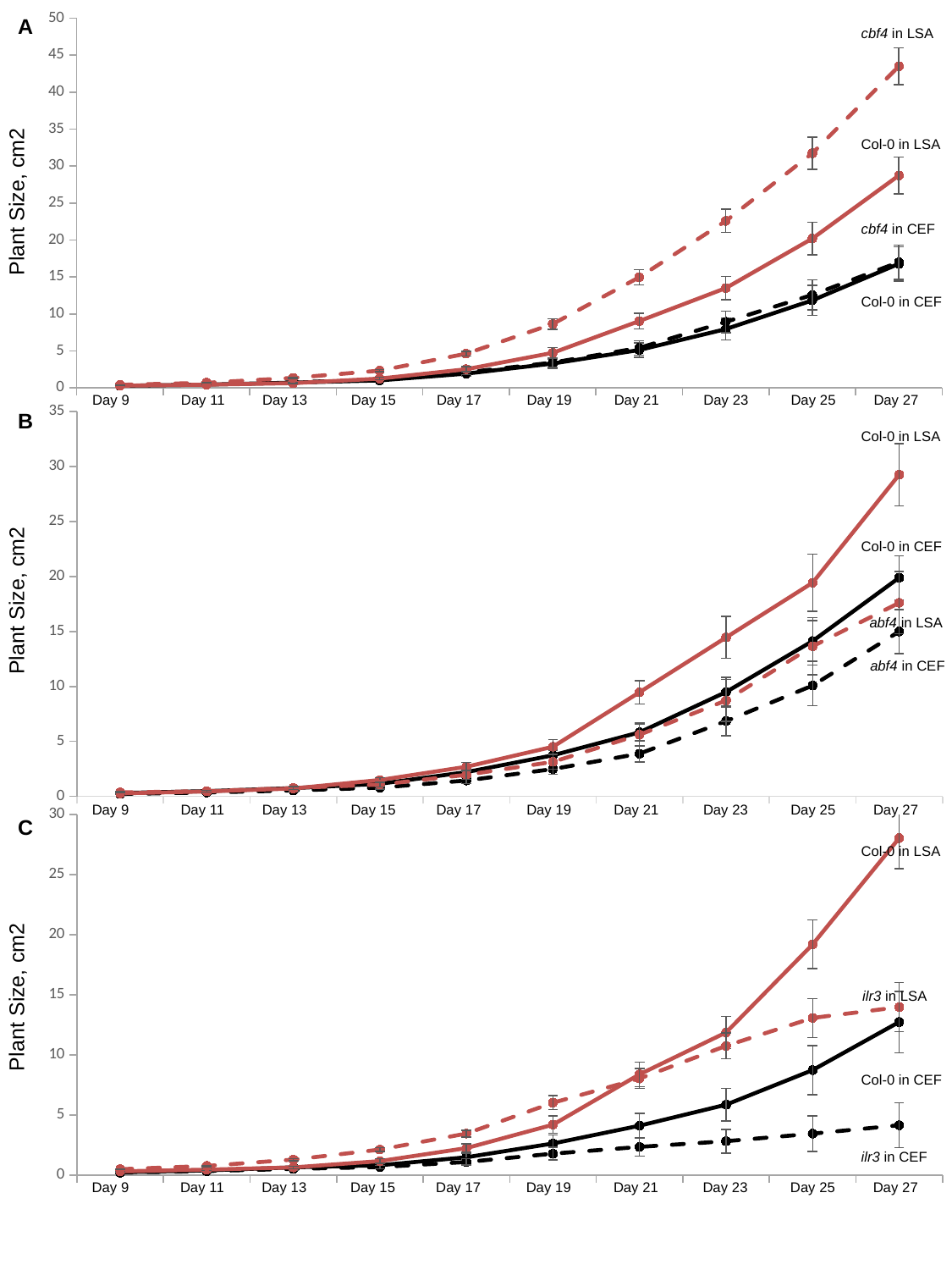

#### Chart
| Category | | | | |
|---|---|---|---|---|
| Day9 | 0.282357142857143 | 0.298642857142857 | 0.294695154886073 | 0.411166666666667 |
| Day11 | 0.445785714285714 | 0.488428571428572 | 0.432 | 0.694583333333333 |
| Day13 | 0.7135 | 0.756142857142857 | 0.63575 | 1.34725 |
| Day15 | 1.0125 | 1.12757142857143 | 1.25458333333333 | 2.32891666666667 |
| Day17 | 1.92192857142857 | 2.09057142857143 | 2.51975 | 4.62175 |
| Day19 | 3.29028571428571 | 3.41585714285714 | 4.73233333333333 | 8.63341666666666 |
| Day21 | 5.12 | 5.40414285714284 | 9.02608333333333 | 14.9561666666667 |
| Day23 | 7.94478571428571 | 8.92907142857143 | 13.49375 | 22.5686666666667 |
| Day25 | 11.8351428571429 | 12.57 | 20.1895 | 31.7244166666667 |
| Day27 | 16.7735 | 17.0000714285714 | 28.72675 | 43.4911666666667 |A
cbf4 in LSA
Plant Size, cm2
Col-0 in LSA
cbf4 in CEF
Col-0 in CEF
Day 9
Day 11
Day 13
Day 15
Day 17
Day 19
Day 21
Day 23
Day 25
Day 27
#### Chart
| Category | | | | |
|---|---|---|---|---|
| Day9 | 0.308625 | 0.214 | 0.272 | 0.36325 |
| Day11 | 0.475125 | 0.353375 | 0.4475 | 0.47575 |
| Day13 | 0.74775 | 0.540750000000001 | 0.71425 | 0.76475 |
| Day15 | 1.14475 | 0.794375 | 1.491 | 1.071 |
| Day17 | 2.20625 | 1.44425 | 2.6935 | 1.96275 |
| Day19 | 3.734 | 2.483875 | 4.512 | 3.14125 |
| Day21 | 5.821875 | 3.885 | 9.47175 | 5.607 |
| Day23 | 9.4935 | 6.84712500000001 | 14.45725 | 8.72425 |
| Day25 | 14.128875 | 10.082125 | 19.43775 | 13.65325 |
| Day27 | 19.893125 | 14.99625 | 29.265 | 17.611 |B
Col-0 in LSA
Plant Size, cm2
Col-0 in CEF
abf4 in LSA
abf4 in CEF
Day 9
Day 11
Day 13
Day 15
Day 17
Day 19
Day 21
Day 23
Day 25
Day 27
#### Chart
| Category | | | | |
|---|---|---|---|---|
| Day9 | 0.235 | 0.207076923076923 | 0.313333333333333 | 0.493545454545455 |
| Day11 | 0.392571428571428 | 0.338076923076924 | 0.455285714285714 | 0.762363636363636 |
| Day13 | 0.648285714285715 | 0.518037275330615 | 0.642571428571429 | 1.28981388091254 |
| Day15 | 0.820857142857143 | 0.671076923076922 | 1.16542857142857 | 2.12409090909091 |
| Day17 | 1.48514285714286 | 1.09015384615384 | 2.24985714285714 | 3.45936363636364 |
| Day19 | 2.62985714285714 | 1.786 | 4.19642857142857 | 6.02136363636364 |
| Day21 | 4.11271428571429 | 2.348 | 8.386 | 8.03563636363636 |
| Day23 | 5.85742857142857 | 2.82392307692308 | 11.8642857142857 | 10.7560909090909 |
| Day25 | 8.73628571428572 | 3.45053686849939 | 19.1998571428571 | 13.076759020714 |
| Day27 | 12.7378571428571 | 4.1513076923077 | 28.0427142857143 | 13.9777272727273 |C
Col-0 in LSA
Plant Size, cm2
ilr3 in LSA
Col-0 in CEF
ilr3 in CEF
Day 9
Day 11
Day 13
Day 15
Day 17
Day 19
Day 21
Day 23
Day 25
Day 27

### Slide 3
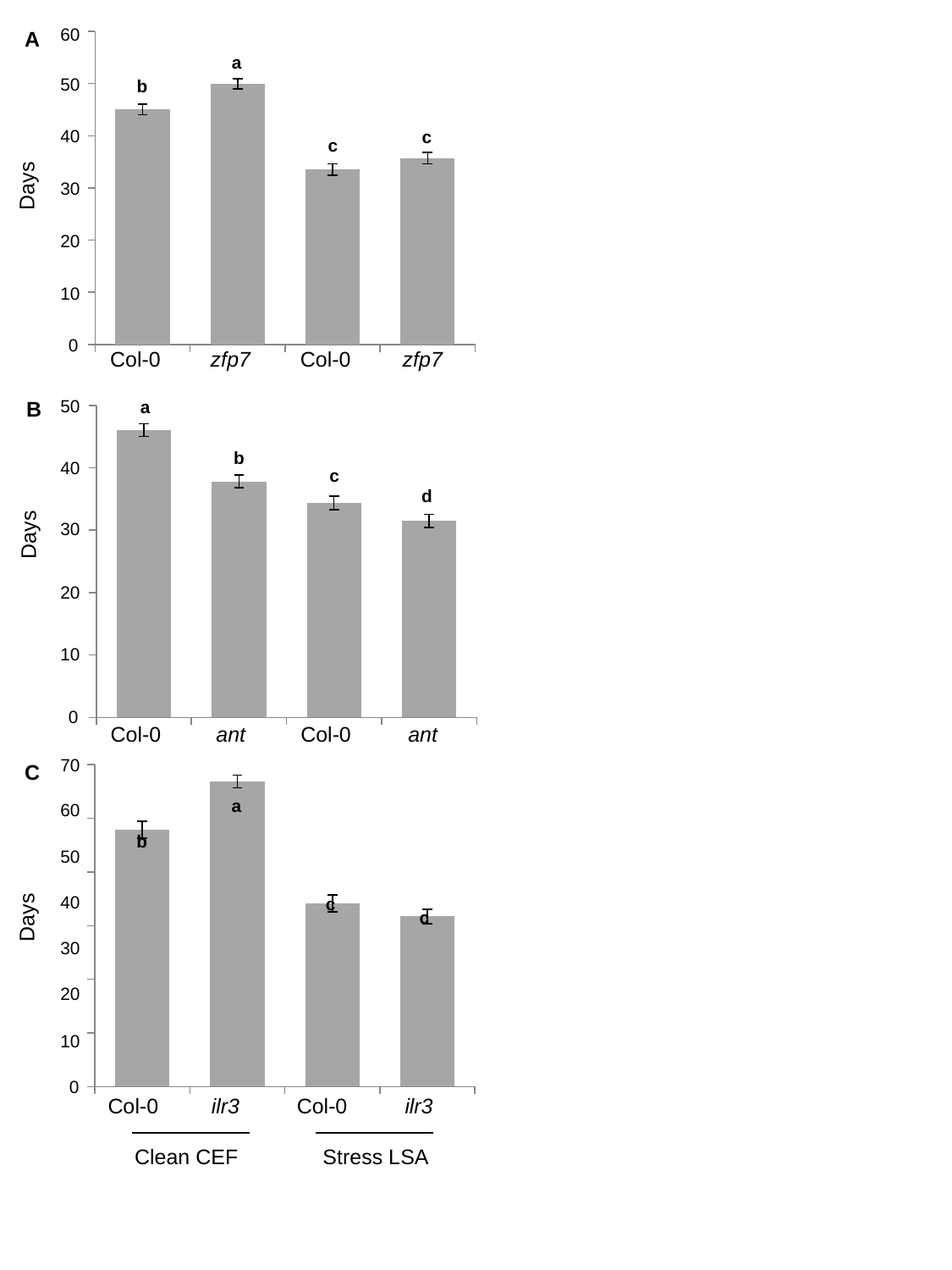

#### Chart
| Category | |
|---|---|60
A
a
50
b
40
c
c
Days
30
20
10
0
Col-0
Col-0
zfp7
zfp7
#### Chart
| Category | |
|---|---|50
a
B
b
40
c
d
Days
30
20
10
0
Col-0
Col-0
ant
ant
#### Chart
| Category | |
|---|---|70
C
a
60
b
50
40
Days
c
c
30
20
10
0
Col-0
Col-0
ilr3
ilr3
Clean CEF
Stress LSA
